# Loci specific epigenetic drug sensitivity

**DOI:** 10.1101/686139

**Authors:** Thanutra Zhang, Anna Pilko, Roy Wollman

**Affiliations:** Institute for Quantitative and Computational Biosciences, University of California, Los Angeles; Departments of Integrative Biology and Physiology and Chemistry and Biochemistry, University of California UCLA

## Abstract

Therapeutic targeting of epigenetic modulators offers a novel approach to the treatment of several diseases including cancer, heart diseases and AIDS. The cellular consequences of chemical compounds that target epigenetic regulators (epi-drugs) are complex. Epi-drugs affect global cellular phenotypes and cause local changes to gene expression due to alteration of a gene chromatin environment. Despite increasing use in the clinic, the mechanisms responsible for cellular changes are unclear. Specifically, to what degree the effects are a result of cell-wide changes or disease related locus specific effects is unknown. Here we developed a platform to systematically and simultaneously investigate the sensitivity of epi-drugs at hundreds of genomic locations by combining DNA barcoding, unique split-pool encoding and single cell expression measurements. Internal controls are used to isolate locus specific effects separately from any global consequences these drugs have. Using this platform we discovered wide-spread loci specific sensitivities to epi-drugs for three distinct epi-drugs that target histone deacetylase, DNA methylation and bromodomain proteins. By leveraging ENCODE data on chromatin modification, we identified features of chromatin environments that are most likely to be affected by epi-drugs. The measurements of loci specific epi-drugs sensitivities will pave the way to the development of targeted therapy for personalized medicine.

## Introduction

The location of a gene on the chromosome is known to affect its expression. Position effect was first observed in *Drosophila* by Muller in 1930 [1,2] and intensively investigated afterward [3–5]. Many years after the original work in *Drosophila*, it is now well documented that gene expression levels are influenced by chromatin environment [2,6–10]. Chromatins play a key role in the regulation of gene expression and are responsible for cell maintenance and differentiation. Chromatin regulation is complex and is an area of active research. The three dimensional structure of the chromatins plays an important role in gene regulation by controlling accessibility of transcriptional machinery and the spatial proximity of a gene from cis regulatory elements such as enhancers. In addition, the specific three dimensional folding will change the spatial distribution of transcription factors and other regulatory molecules such as lncRNAs. The spatial proximity of these regulatory molecules then play a key role in controlling gene expression patterns. The three dimensional structure itself is highly correlated with specific histone and DNA modification patterns. Overall, the complex multi-layered regulation of chromatin on gene expression pattern causes each gene to exist in a unique chromatin environment that plays an important role in determining gene expression distribution, i.e. both its average level as well as population variability [6,9,11].

The proper regulation of gene expression is vital for health and dysregulation of gene expression is associated with a large number of pathologies. Advances in DNA sequencing allow the classification of the specific disease based on the underlying changes of gene expression, the basis of large parts of precision medicine approaches. Given the large knowledge that is accumulating on what changes in gene expression are associated with disease conditions, it is only natural to attempt to correct these pathologies by modification of underlying gene expression patterns. This quest has long history with initial attempts related to antisense oligos[12]. Similarly, the discovery of RNA interference (RNAi) sparked many attempts to develop therapies based on RNAi with the goal of manipulating gene expression [13]. However, despite the conceptual simplicity, translating these concepts into therapy was challenging[14–16].

Given the influence of local chromatin environment on gene expression, strategies that target epigenetic regulators are being investigated. Two main strategies are the pharmacological use of epi-drugs to influence gene expression and targeted approaches for epigenetic editing. Pharmacological approach uses inhibitors to the readers/writers/erasers of epigenetic marks. The pharmacological approach that is being developed to address a wide range of diseases is continuously expanding [17–23]. Multiple targeting strategies for epi-drugs are being explored including specific loss and gain of function [24–31], synthetic lethality [32–35], and to overcome drug resistance [35–37]. A common theme across these strategies is the use of epi-drugs to manipulate gene expression patterns e.g. suppress oncogenes or activate tumor suppressor genes [38]. However, the precision of epi-drugs induced gene expression targeting, i.e. the fraction of overall changes to gene expression that are desired for therapy, is currently very low. This low precision limits the usability of epi-drugs [39–41]. The alternative strategy is based on targeted recruitment of epigenetic modulators into specific sites. CRISPR mediated sequence specific targeting of epigenetic regulators is used to cause changes in gene expression pattern of specific loci. [42–44] The key advantage of epigenetic engineering is their precision. However, many challenges have to be addressed before these approaches can be translated into the clinics.

Despite the popularity of the use of epi-drugs to cause changes in gene expression patterns, there are many unknowns resulting from gaps in existing measurement capabilities of the effect of epi-drugs on gene expression. Epi-drugs change gene expression due to direct, locus-dependent changes, and indirect or nonspecific effects [38]. Existing approaches to identify direct effects of epi-drugs rely on a combination of RNAseq and multiple ChIPseq to show that gene expression changes are coupled to changes in local histone modification [45]. However, the reliance on ChIPseq makes this approach limited as these measurements are only semi-quantitative [46], it is often hard to interpret their functional effects on gene expression [47], and they are challenging to scale to large numbers of samples [48]. Therefore, it is currently impossible to rigorously identify changes to gene expression that are due to specific modifications to the chromatin environment and not a result of non-specific or indirect effects. Therefore, there is a gap in current capabilities of mapping the specific and local impact of drug manipulating chromatin modifiers on gene expression.

Here we developed new measurement technology for the Massive And Parallel Measurement of Epigenetic Drug Sensitivity (MAPMEDS). MAPMEDS is based on comparison of drug effect at specific locus compared to the drug effect of hundreds of other loci. Statistical comparison of the drug effects on expression of a reporter fluorescent protein allows the deconvolution of global effects and locus specific sensitivities. MAPMEDS utilizes DNA barcodes to pool together the time consuming step of genomic position identification of the reporters. Split-pool approach enables mapping of DNA barcodes to individual reporter cell lines. Using MAPMEDS we demonstrate the widespread existence of loci specific sensitivities for three epi-drugs that target histones acetylation, DNA methylation and proteins with bromodomains. By leveraging ENCODE data on the chromatin environment in each location, we show what types of environments are more susceptible to the different epi-drugs. Overall, these results shed light on how epi-drugs cause changes in gene expression, the information that can be used for development of more precise targeting strategies.

## Result

### Overview of MAPMEDS

MAPMEDS is based on two key innovations: 1) A split-pool strategy for the creation of cell lines that uses DNA barcodes to identify what DNA labeling each cell line has and what is the genomic integration position of that barcode (Fig. 1a-d). Each expression reporter incorporates unique DNA barcode into reporter cassette that contain identical promoter and fluorescent reporter. These barcodes serve as unique identifiers for mapping the genomic locations of reporters and revealing clonal identities without the need of individual genomic extraction. The use of split-pool encoding allows the mapping between isoclonal line expression and the DNA barcode and thereby connecting genomic information with expression measurements. 2) A new strategy that uses in-well controls and statistical tools enables the separation of gene expression changes into locus-specific and global non-specific (Fig. 1ef). The use of in-well controls minimizes batch effects and is a key to identify locus specific sensitivities. Collectively, these two steps allow the parallel creation of large number of reporter cell lines and their use to identify loci specific epigenetic drug sensitivities.

**Figure 1:**
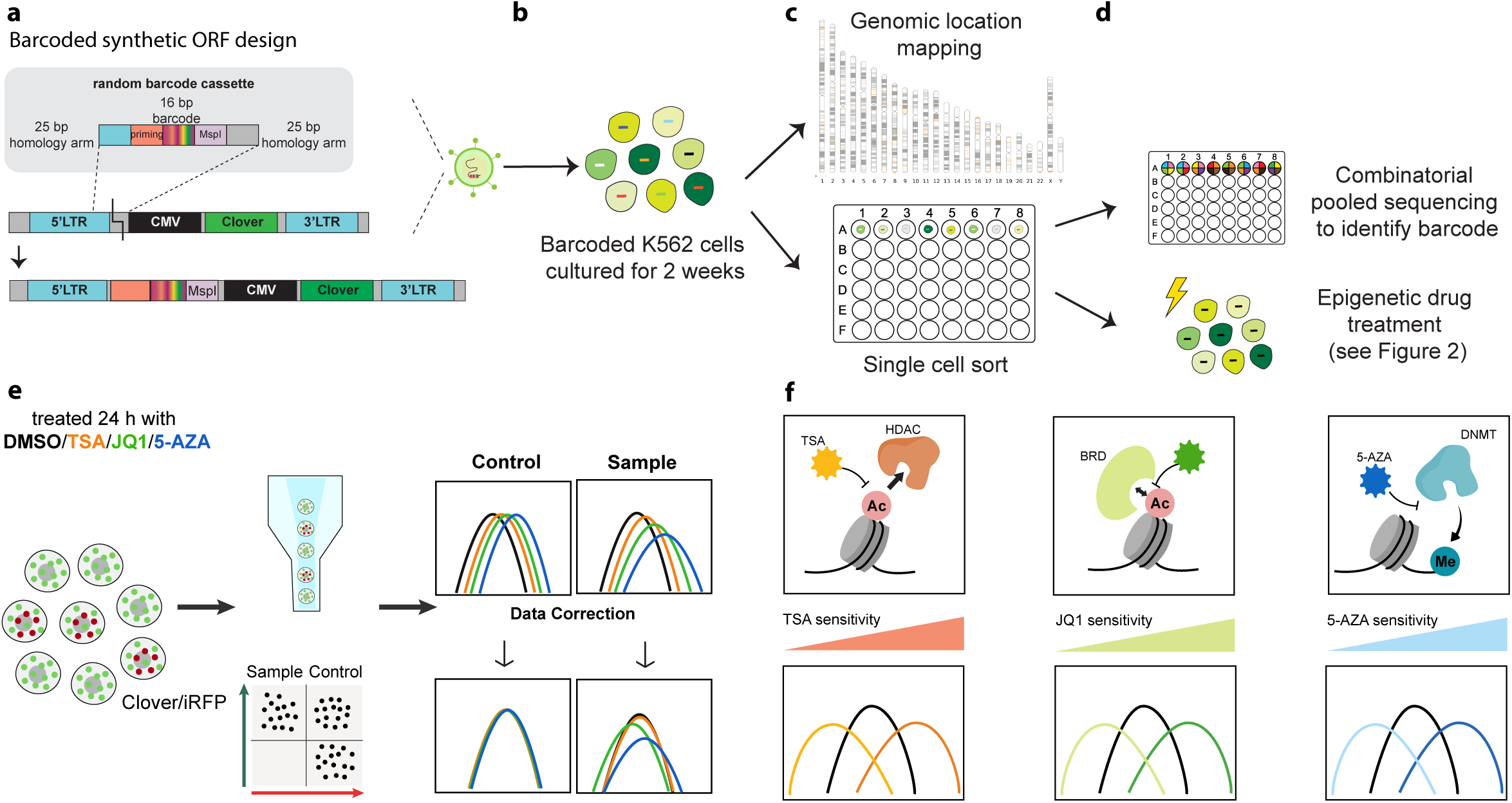
An overview of MAPMEDS. **(a)** Schematic structure of barcoded lentiviral constructs. The library vector contains short barcode, CMV promoter and mClover fluorescent protein as a reporter. The barcode is random 16-bp-long DNA with repeats of A,G and T. MspI restriction site is integrated upstream CMV promoter for genomic location mapping. **(b)** Barcoded lentivirus was packed and transduced into K562 cells at low MOI to create founder cells with singly integrated reporter. **(c-d)** Barcoded founder cells, selected by flow cytometry, were expanded for two weeks and split into two pools. Cells in the first pool were collected for locating reporter integration site. Founder cells in the second pool were sorted into 96-well plates to establish clonal cell lines. Barcode of each clone was simultaneously identified by split-pool encoding and deep sequencing. Library of characterized reporter clones is a useful resource to examine loci specific epigenetic drug sensitivity. **(e)** Loci specific effects were decoupled from global effects through mixing individual barcoded clones of interests with control cells expressing mClover and IRFP from multiple integration sites. Co-cultured cells were treated with TSA, JQ1 and 5’AZA for 24 hours. Expression of reporter proteins were measured by flow cytometers. Distribution of mClover expression in control cells was used to remove global effects of drugs. **(f)** A cartoon illustrating known mechanisms of actions of Trichostatin A, JQ1 and 5-Azacytidine.

MAPMEDS utilizes DNA barcoded expression reporters integrated as single copy per cell. In order to generate founder cells with one copy of barcoded reporter, lentiviral transduction at low multiplicity of infection (MOI) was used. In short, we first created a library of lentiviral plasmids containing barcode, MspI restriction site, cytomegalovirus (CMV) promoter and GFP variant, mClover [49] (Fig.1a). The barcodes are 16-bp sequences of random Adenosine, Thymine and Guanine. Cytosine is excluded to keep barcode intact for future application to examine DNA methylation through bisulfite conversion. Barcoded lentiviruses were packed and transduced at MOI of ∼0.01 into K562 cells, leukemia cell line with abundant available epigenetic profiles [50] (Fig.1b). Reporter K562 cells were selected at 72 hours post-transduction by fluorescent activated cell sorter (FACS) to establish a pool of 3,000 founder cells. Founder cells were expanded for 2 weeks and split into two pools (Fig.1c). One half was used to identify the genomic location using inverse PCR [6]. The other half was used to establish individual isoclonal lines through single cell sorting. Once the cell lines were established, we utilized a combinatorial pooling approach to combine all the cell lines into a small number of pooled samples that were used to identify the barcode identity, and hence the genomic integration site, of the ORFs for all cell lines (Fig. 1d). The result of this procedure is a library of clonal cell lines that can be used as a powerful resource to examine drug sensitivity at diverse epigenetic environments.

In order to isolate locus-specific changes from any global changes in gene expression patterns, we implemented a new strategy using in-well controls and statistical tools. A reference “non-specific” cell population was created by using a polyclonal population of multiple integration cells so that the overall population has thousands of integration sites of same reporter cassettes. Far-red fluorescent protein iRFP670 [57] was used to mark this reference population. We split these control cells and co-cultured with each target cell line in multiwell plates.

### Integration landscapes of reporters

To map the integration sites of reporters, we split half of founder cells into three sub-pools and further expanded the population. The first pool was used to reveal a list of genuine barcodes. We detected 756 candidate genuine barcodes after two weeks of culture. Two other sub-pools are technical replicates for locating reporter integration sites by an inverse PCR method coupled to paired-end high-throughput sequencing (Fig. 2a). In short, genomic DNA from each pool was isolated, digested with MspI enzyme and self-ligated. Barcodes and adjacent genomic DNA were amplified and deep sequenced. After barcode demultiplexing, genomic sequence was mapped to human genome assembly GRCh38. Genomic coordinate was assigned to corresponding barcode when mapped results from two replicates are matched. We observed reporter integration throughput the genome (Fig.2b, Fig.S1a) with enriched pattern similar to previous study (Fig.S1b) [8]. Reporters were integrated at various genomic environments serving as diversified resources to study epigenetic drug sensitivity.

**Figure 2:**
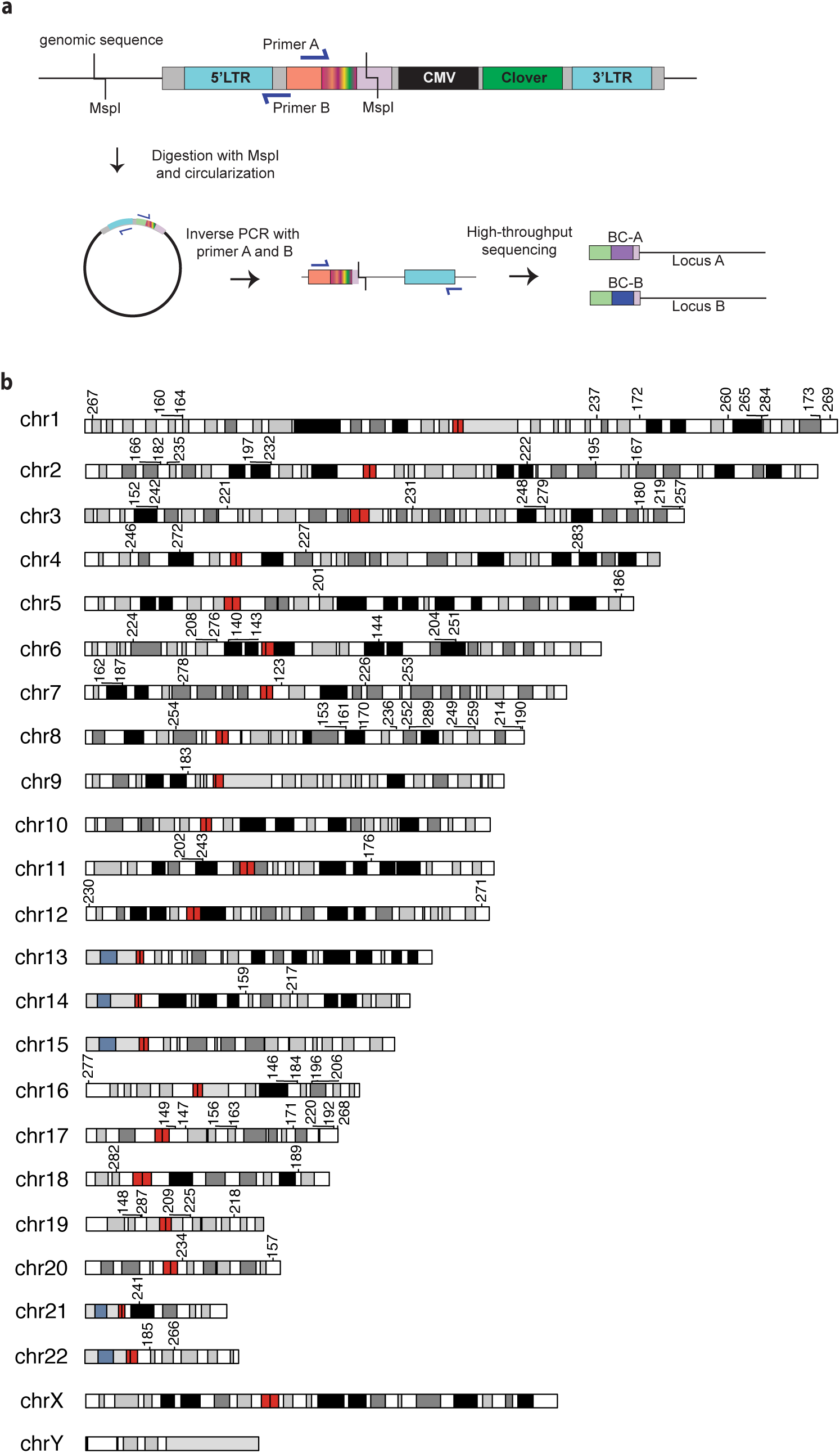
Diverse insertion landscapes of barcoded reporter. **(a)** Reporter mapping by inverse PCR. Genomic DNA of founder cells was extracted, digested with restriction enzyme MspI and self-ligated to stitch barcode with its neighboring genome. Ligated product was amplified and followed by next generation sequencing. **(b)** Ideogram plot displaying reporter integration sites of individual clones in the library. Centromere position is indicated in red and stalk is marked in light blue. Heterochromatic region, which tend to be rich with adenine and thymine and relatively gene-poor, is represented by black and variation of grey. R-band in white on the ideogram is less condensed chromatin that is transcriptionally more active.

### Scalability and Robustness of MAPMEDS

Pooled-sample sequencing is a cost-effective and practical strategy for many studies, especially the ones related to the discovery of rare mutation and single nucleotide polymorphism associated diseases [51–53]. Combinatorial pooled sequencing is an extension of standard pooled sampling where each sample exists in a few pools creating a many to many mapping between experimental conditions and samples. Combinatorial pooling improves the sensitivity and robustness of pooled sequencing since it includes built in error corrections. It reduces the cost and time for library preparation exponentially [54–56].

Conceptually, in combinatorial pooled sequencing, the identity of each sample is encoded in the composition of pools and this pooling pattern serves as a reference for decoding sequences belonging to corresponding sample (Fig.3a). Clonal lines are mixed into few pools according to a predefined design. Each individual pool is genomic DNA extracted, barcode amplified and index tagged using nested PCR. Amplicons from all pools are mixed together and sequenced. Subsequently, sequencing data is sorted by barcodes and their appearance patterns determined by the well indexes. The measured patterns are compared with the design pooling signatures to match barcodes with clone numbers.

**Figure 3:**
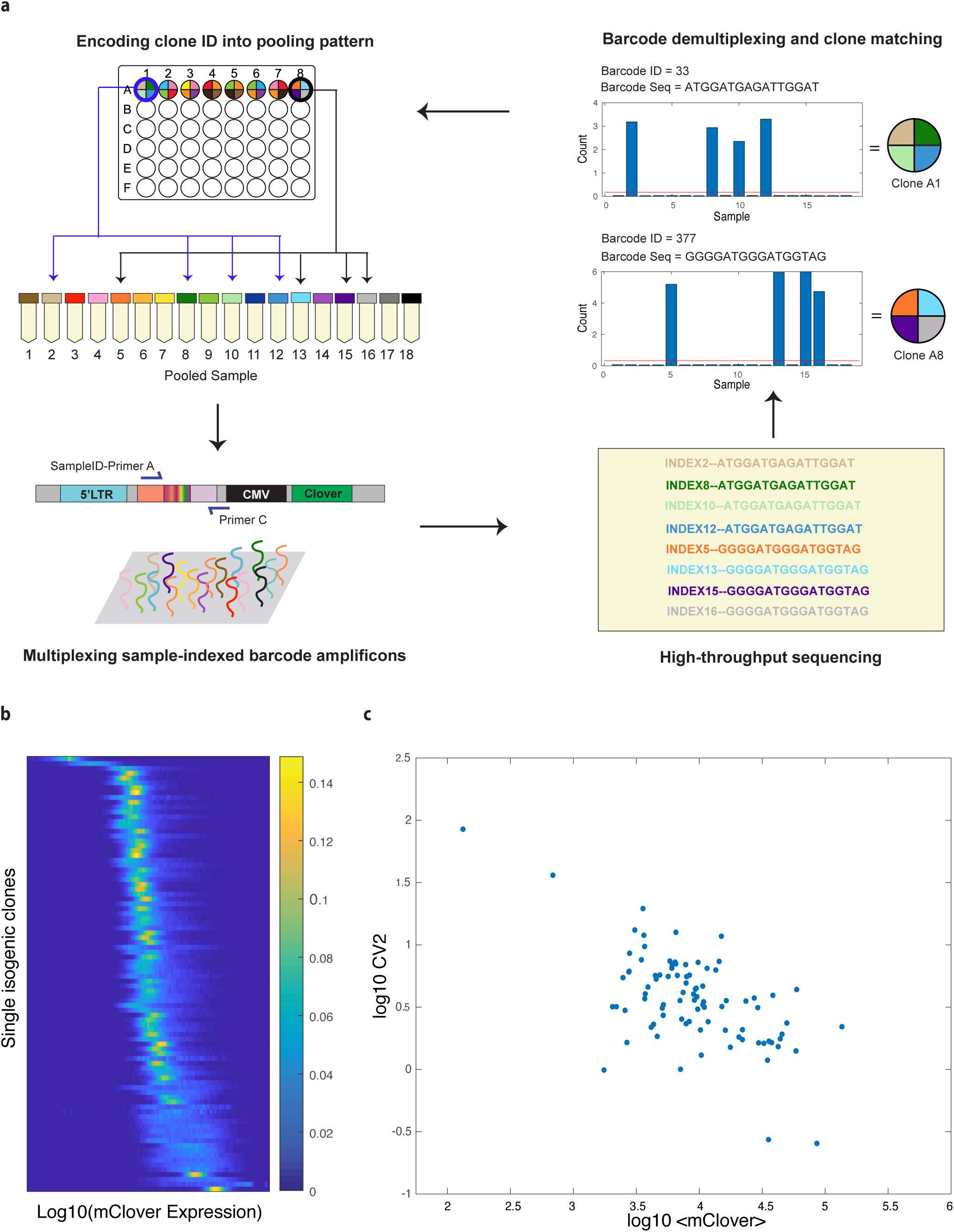
Combinatorial pooling massively and parallely identify barcode of individual clones. **(a)** The combinatorial pooled sequencing involves encoding and decoding steps. Each individual clone was split into 4 out of 18 pooled samples according to the designs. Genomic DNA from each pooled sample was extracted and barcodes were amplified and labeled with 6-bp sample index. Amplicons from all samples were mixed together and prepared for NGS. Detected barcodes were deconvoluted to match barcode ID with clone ID. **(b)** The expression distribution of mClover protein was measured by high-throughput flow cytometer and displayed by stacked probability density function. Each row in the heatmap represents a single histogram from a single clonal line with the probability density function colorcoded. **(c)** A scatter plot of reporter expression noise, measured as log-transformed squared coefficient of variation (CV^2^), and mClover mean across all examined positions affirms unique chromatin environments across the genome.

The use of combinatorial pooling circumvents the need for individual genomic extraction and PCR per clones which significantly reduces the cost of reagents and hands-on time for sequencing preparation. In our design, each clone is mapped into 4 selected pools from total of 18 pools and this set-up allows up to 18-choose-4 or 3060 sample identification. This approach is highly scalable because sample size can be easily increased by adjusting number of pool and bit of code. We checked the accuracy of barcode identification using combinatorial pool sequencing by targeted PCR and Sanger sequencing. For 5 randomly chosen clones, the barcode deconvolution was all corrected which confirm the robustness of our method.

### Chromosomal Position Effects on Protein Expression Distribution

To support the notion that each integration site exists in a different chromatin environment, we examined the positional effect of our expression reporters. Across nearly a hundred positions, we observed variable expression distribution of mClover protein (Fig.3b). Some clones are completely silent while many are highly expressed with expression average of approximately 1000-fold higher than the lowest-expressing cells (Fig.3c). This variation range is comparable to a study that measured averaged mRNA expression across mouse genome [6], but higher than other studies carried out in bacteria or yeast [7,9,10]. Broader magnitude of positional effect in expression level detected in mammalian genomes may come from their large and complex genome organization and the discrepancy in experimental design and techniques.

Our observation on variable expression distribution confirms that the location where reporter gene inserted affects its expression and differences in genomic landscapes and epigenetic profiles are suggested to explain such differential expression in several studies. For example, lamina-associated domain and chromatin compaction significantly attenuates transcriptional activity [6] and certain histone marks, including H3K36me3, are correlated with expression level of the reporters [9,10]. Intuitively, epigenetic drugs that target chromatin regulator should also be impacted by distinct genomic environments. However, such loci specific sensitivity has not been previously measured and this motivates us to systematically measure positional effects on the sensitivity of epigenetic drug using our library of isogenic clones established and characterized by MAPMED.

### Epigenetic drugs show position-dependent sensitivity

As a proof of concept, we chose three epigenetic drugs representing three mechanisms of inhibition. Trichostatin A (TSA) is histone deacetylases (HDACs) inhibitor [58,59] and effective in the treatment of several types of cancer including promyelocytic leukemia [60], lung cancer [61], breast cancer [62] and also enhancing the response of chemotherapy of multiple cancers [63,64]. JQ1 is a small-molecule inhibitors of BRD2 and BRD4, members of the bromodomain and extra-terminal domain (BET) protein family. JQ1 competitively blocks the binding of bromodomain proteins and acetylated chromatin which results in transcriptional attenuation [65,66]. JQ1 has been reported as a promising cancer therapeutic strategy in several cancers [67–69]. Azacytidine is a cytidine analogue that inhibits DNA methylation through loss of DNA methyltransferase (DNMT) activity [70]. Azacitidine was approved by the U.S. Food and Drug Administration (FDA) for the treatment of all subtypes of myelodysplastic syndrome (MDS) since 2004 [71]. These three drugs were added to co-cultured cells and incubated for 24 hours before sample collection and fluorescent quantification.

The expression level of mClover and iRFP was measured using high throughput sampler (HTS) flow cytometer. Cells from each sample were separated into reference and target cells based on iRFP gating. To quantify the site specific effect, we normalized the expression levels of the sample cells by the average change between the drug and DMSO in the control population. After correction, the site specific effect of a drug that are not captured by reference population were quantified by the Kolmogorov–Smirnov (KS) statistics that compares distribution similarity between sample cells under drug and sample cells with DMSO (Fig 4a). Any effects that are only site specific are effectively measured as changes beyond those also occur in the reference polyclonal population. Differential magnitudes of drug sensitivity were observed across examined locations. Some loci, such as clone 224 and clone 260, show indifferent distribution of mClover in all conditions, but many positions are either hyposensitive or hypersensitive to at least one of epigenetic drugs (Fig.4b). These non-uniform responses indicate unique chromatin environment at each genomic location.

**Figure 4:**
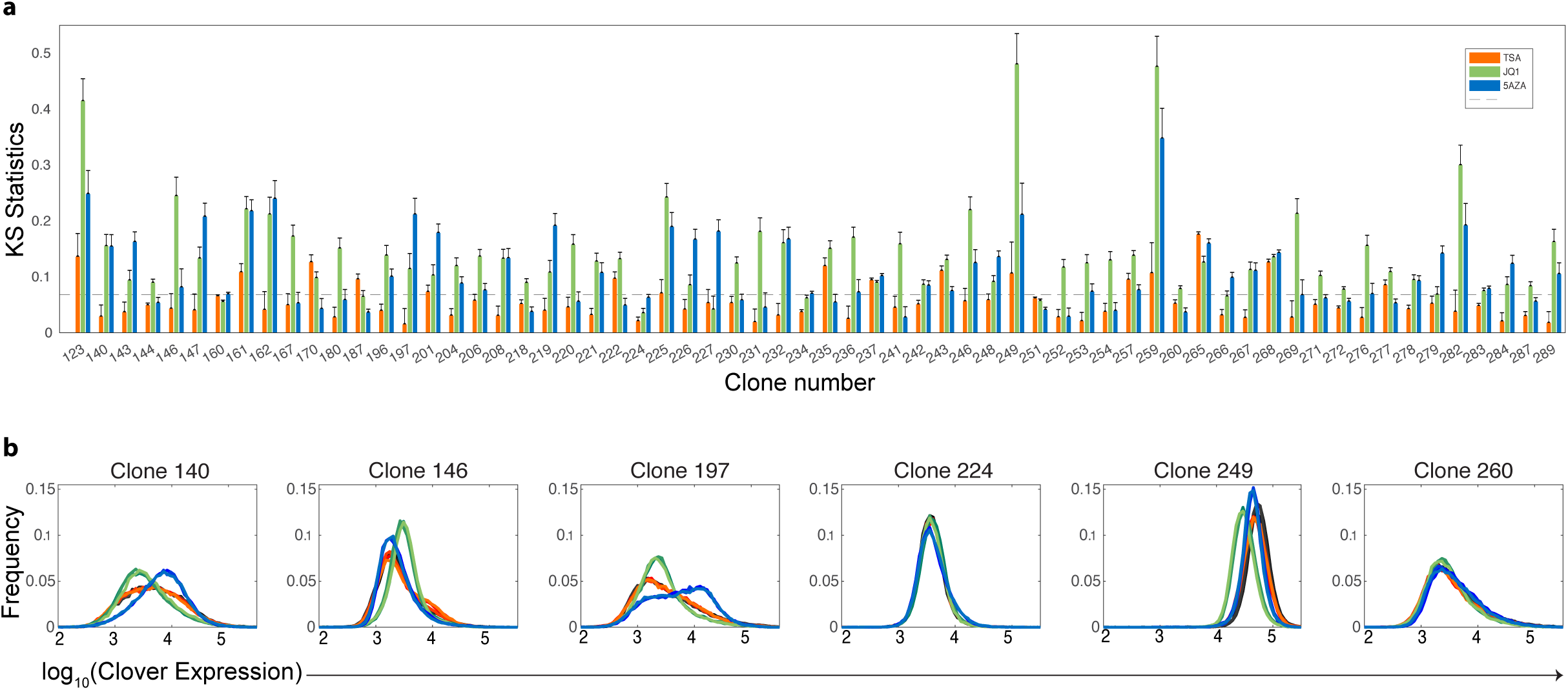
Chromosomal position effects influence magnitudes of epigenetic drug effects. **(a)** A bar graph of calculated the test statistic of two-sample Kolmogorov–Smirnov test between epigenetic drug treatment and DMSO. **(b)** Histogram plot showing mClover distributions of selected clones after 24-hour treatment of DMSO(Black), TSA(Orange), JQ1(Green) and 5 Azacytidine(Blue).

The distributions of KS statistics in control cells fall within the three standard deviation limits. (Fig. S2). Therefore, we used this criteria to identify the number of ‘hits’ per drug as it provides a non-parametric estimate of our false discovery rate to be <0.015. Our drug screening shows ∼16% and ∼13% of the positions that have down-regulated and up-regulated reporter expression after TSA treatment. The number of hits in TSA screen is lower than those in JQ1 and 5-Azacytidine. The percentage of down-regulated and up-regulated loci are 64% and 30% in JQ1 and 40.6% and 29.7% in 5-Azacytidine respectively. Smaller fraction of TSA-sensitive sites observed in our data is in agreement with other studies findings that HDAC inhibitors only target less than 10% of the genome [72–74]. Overall, our data demonstrates a chromosomal position effects on epigenetic drug sensitivity.

### Analysis of histone modification profiles identifies chromosomal environments susceptible to JQ1 sensitivity

We compared our maps of epigenetic drug sensitivity to a collection of available epigenomic map from K562 cells focusing on histone modifications. ChIP-seq signals, expressed by fold change over control, in a window of 10 kb around barcodes of interest were considered. We note that histone modification profiles were mapped in K562 cells without any genetic engineering. Insertion of reporters potentially change the pre-existing epigenetic landscape or cause sequence-specific or protein-specific interactions between regional chromatin and synthetic ORF. However, previous studies suggest that integrated reporters generally do not perturb the chromatin landscape but adopt the local chromatin state [10,75]. Complex sequence- or protein-specific interactions between local chromatin and synthetic reporters were not observed in previous study as well [11].

After selecting top JQ1-sensitive clones with distinct downregulation and upregulation of reporter expression (Fig.5a), we compared their epigenetic profiles. Interestingly, we found JQ1 sensitivity is associated with certain histone modifications (Fig.5b). Structural study reveals the binding of JQ1 to the acetyl-lysine binding pocket of BET bromodomains [65]. Such competitive binding disrupt bromodomain/acetyl histone interaction and therefore transcriptional activation. Genomic regions enriched in acetylated histones were hypothesized to display higher magnitude of reporter downregulation. Indeed, we observed that clones with lower expression after JQ1 treatment show higher enrichment of H3K9ac and H3K27ac. Moreover, we also found that these clones have significantly higher enrichment of H2A.Z. This data supports previous studies demonstrating that BRD2 interacts with H2A.Z to mediate transcription initiation [76,77] and thus suggests that our assay recapitulated well-established results.

**Figure 5:**
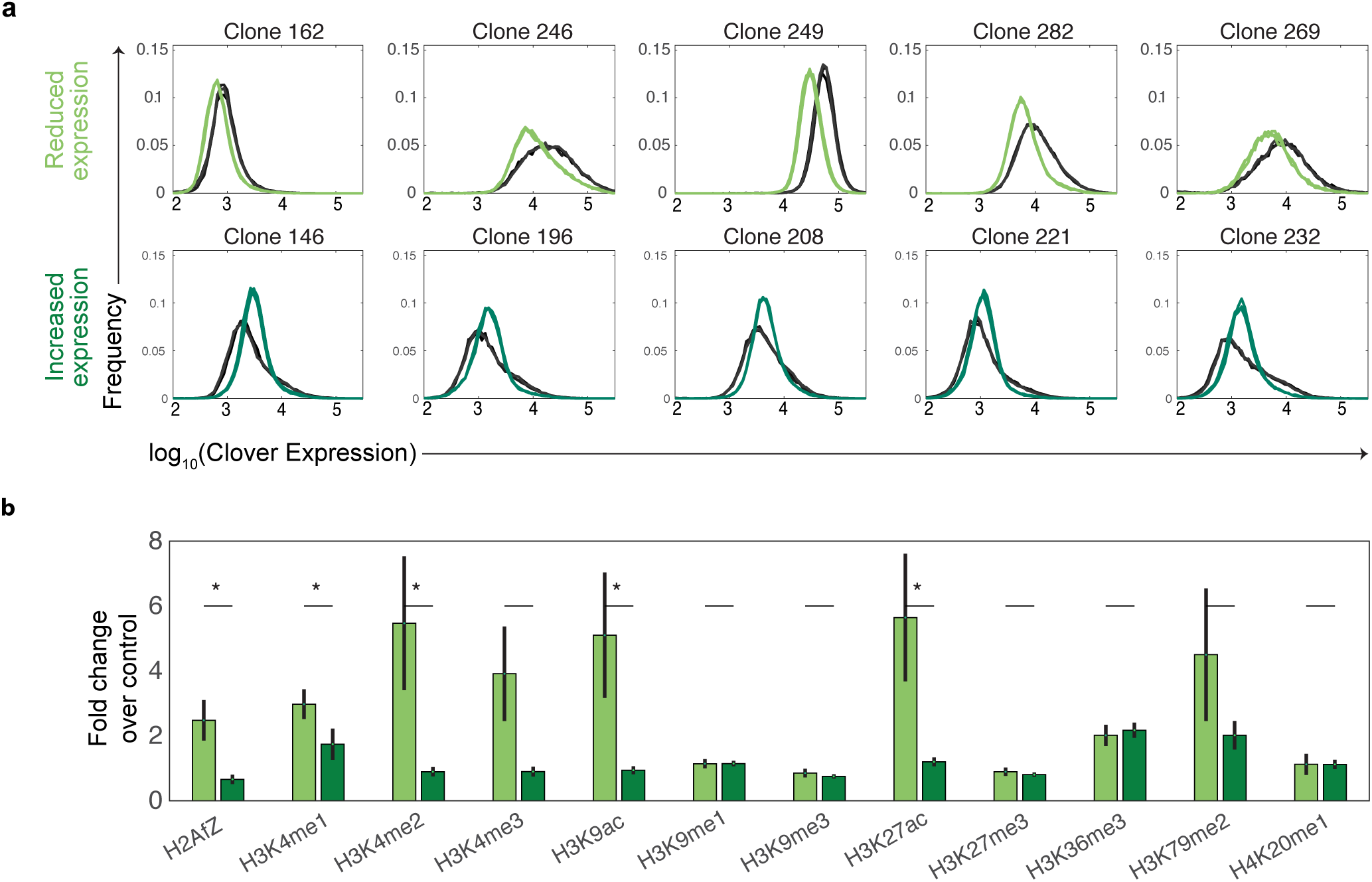
H2A.Z influences sensitivity of bromodomain inhibitor to expression alteration. **(a)** Examples of mClover distribution from clones showing significant reduced (top row) and increased (bottom row) mClover expression to JQ1 drug. **(b)** Bar graph shows histone enrichments for comparison of positions displaying reporter down-regulation (light green) versus those exhibiting up-regulation (dark green) after JQ1 treatment. Fold change over control of each histone mark was averaged within a window of 10 kb. The p-values were determined by two-sample t-test.

### Differential sensitivity to 5-AZA treatment happened through DNA methylation-independent mechanism

Considering Azacytidine as a well-known inhibitor of DNA methylation, it is likely that genomic regions with hypermethylated promoter will response to such epigenetic drug the most. Moreover, CMV promoter used in our reporter is highly enriched in CpG sites which should be susceptible to DNA methylation. Therefore, we hypothesized that reporters inserted at diverse genomic environments will have different DNA methylation level and result in differential drug sensitivity. To test this hypothesis, we examined DNA methylation at CMV promoter, which contain 30 CG sites, using target bisulfite sequencing (Fig6a).

**Figure 6:**
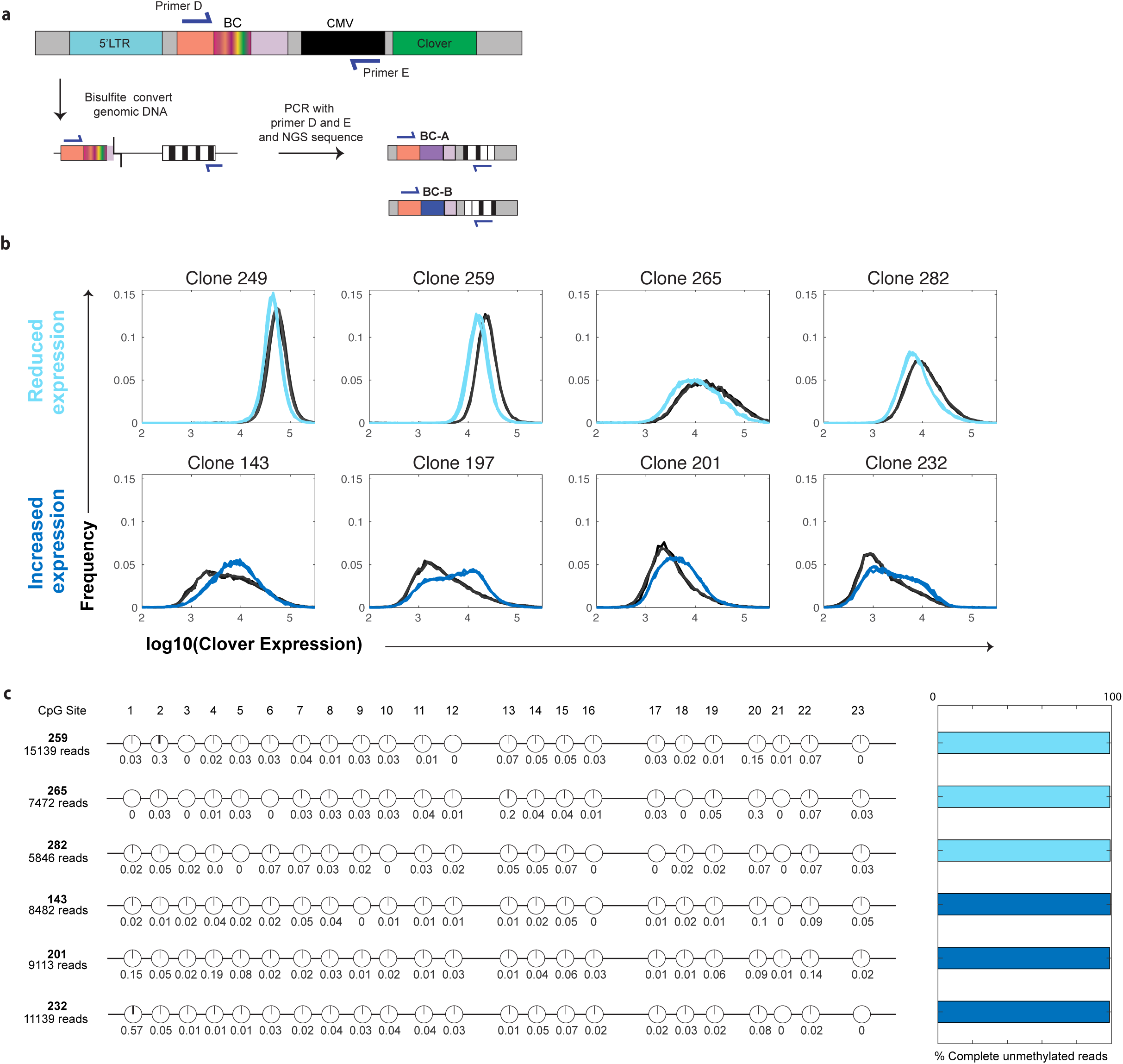
Chromosomal position effects 5 Azacytidine sensitivity through DNA methylation-independent mechanism. **(a)** DNA methylation profiles of selected clones were simultaneously revealed by bisulfite conversion, targeted PCR and deep sequencing. **(b)** Examples of mClover distribution from clones with reduced expression (top row) and induced (bottom row) to 5 Azacytidine. **(c)** Locus-specific bisulfite sequencing of the CMV promoters of representative clones exhibiting hyposensitivity and hypersensitivity to 5 Azacytidine.

Surprisingly, even in cases were we observed distinct reporter downregulation and upregulation in two groups of top sensitive clones (Fig6b), their CMV promoters are mostly unmethylated and indistinguishable (Fig6c). Our data suggests that differential sensitivity to 5-Azacytidine treatment may not result from the direct inhibition of promoter methylation but instead reflect changes in cell regulatory machinery that has stronger effects on these loci than average. This observation is in agreement with previous studies that report 50-60% of induced genes by 5-Aza-CdR did not have CpG islands within their 5’region [78,79]

## Discussion

Here we developed a new method MAPMEDS that can identify loci specific drug sensitivity in a robust and scalable manner. The method uses DNA barcoding to generate cell lines with known integration position of an expression reporter at genome scale and includes a statistical procedure to quantify the locus specific effect that a drug has on gene expression. We used MAPMEDS to evaluate the position specific effects of three common epi-drugs. We found that up to 60% of positions change in a manner that is different than average including both hyper and hypo sensitivity. Through analysis of the chromatin features that are enriched in sites with hyper/hypo drug sensitivities we were able to characterize what aspects of chromatin environment make it more (or less) sensitive to a specific drug. The development of MAPMEDS have both translational and basic science implications.

Locus-specific sensitivity measurements will support the development of new treatment strategies that use existing drugs and the development of new, more precise drugs. Identification of correct dosage of epi-drugs is challenging [80–82]. Comparison of the dose-response curve of the overall change in gene expression to the dose-response changes in gene expression that are due to locus-specific modification will help identifying drug concentrations that maximally impact target genes while limiting non-specific effects. Similarly, it will be possible to improve the precision of drug combinations [19,83]. New drugs and lead compounds could be identified based on predefined desired locus-specific changes in expression patterns. For example, in breast cancer that develops resistance to PI3K therapy, it was shown that co-treatment with a bromodomain inhibitor JQ-1 helps mitigate drug resistance since it silences the compensatory upregulation of RTKs [84]. The use of JQ-1 to achieve such desired effect is limited by the fact that JQ-1 has very broad effects on expression changes across the genome. The tools we propose to develop will allow screening for new compounds and JQ-1 derivatives that maximizes the effects on the desired genes such as EGFR and INSR while limiting other undesirable changes.

Precision medicine is based on stratification of patients and assigning specific therapies based on the molecular information often associated with the key aberrant pathways. Therefore, in many cases, the nature of needed changes in gene expression are known. Treatment is limited due to the lack of therapies that can cause such desired changes in gene expression. Current manipulations of gene expression patterns using epigenetic targeting drugs such as JQ-1 are imprecise[39,40]. The new measurement technology developed here will support the future development of more precise epigenetic drug-mediated gene expression targeting. The reduction in side effects and the ability to screen for locus-specific changes in gene expression will lead to a large array of new therapies.

From basic science perspective, our understanding of chromatin regulation of gene expression is far from complete. MAPMEDs has the capacity to generate large datasets that look at changes across many positions and epigenetic drugs. Such large dataset will provide key insights into how changes in the local chromatin environment can affect gene expression in a manner isolated from any global changes. The use of DNA barcodes as part of MAPMEDS enables the pooled measurement of changes to local chromatin environment at scale. These data will provide invaluable insight into chromatin regulation.

MAPMEDS have two unique aspects that set it apart from other measurement approaches such as TRIP and BHIV [6,8] that aimed at measuring positional effects at scale. Unlike other approaches, MAPMEDS is based on the library creation of cell lines. The use of library of cell lines provides single cell data on changes in expression variability. Indeed, many of the locus specific drug effects we saw did not simply shifted the population but changes the shape of the distribution. These effects would have been missed with bulk population measurements. Additionally, once the initial work in creating the cell line library was invested, the measurement of the drug effects are straightforward and can be scaled to large drug libraries. Other approaches such as TRIP and BHIV will require full RNA sequencing for each drug tested. The down side of the cell line library approach is that it is hard to scale it to more than a few hundred sites. However, recent advances in in-cell barcodes make it possible to generate the cell line library in pooled format, opening the way to a few orders of magnitude increase in the number of positions that can be measured. Future development of MAPMEDS to include these in-cell barcodes will further increase it’s utility as a platform for the discovery of more precise and useful epi-drugs.

## Acknowledgements

We are thankful to Robert Foreman for his help with mapping of reporter integration sites. This work was funded by NIH grants to RW EY024960 and GM111404. T.Z. was also supported by Thailand Her Royal Highness Princess Maha Chakri Sirindhorn fellowship.

## Author Contributions

TZ and RW conceptualized the experiments and data analysis.TZ performed the experiments and performed data analysis. TZ and RW wrote and edited the paper.

## Declaration of Interests

The authors declare no competing interests.

## Methods

### Cell Lines and cell culture

The human K562 cells(Sigma-Aldrich) were grown at 37 °C in RPMI 1640 medium (Gibco) supplemented with 10% FBS (Gibco), 1% penicillin-streptomycin (Gibco) and 1% GlutaMAX (100x) (Gibco) under a 95% air and 5% CO2 atmosphere.

### Construction of library reporter plasmid

The based plasmid without barcode was first constructed to contain the following elements. Lentiviral production units include HIV-1 truncated 5’ LTR, HIV-1 packaging signal, HIV-1 Rev response element (RRE), HIV-1 truncated 3’ LTR and Central polypurine tract (cPPT). These components allow proper viral packaging and viral integration into host cells. As a transcription unit, we used cytomegalovirus promoter (CMV) to drive expression of the reporter gene encoding yellow-green fluorescent protein (mClover). Woodchuck hepatitis virus posttranscriptional regulatory element (WPRE) is placed after mClover to enhances mRNA stability and protein yield. Ampicillin resistance gene (β-lactamase) is included for selection of plasmid in bacterial cells.

To generate barcoded plasmid libraries, based lentiviral plasmid was cut upstream of the CMV promoter by ClaI restriction enzyme and purified by ethanol precipitation. The inserted cassette of 127-bp-long oligonucleotide containing a random 16-bp-long barcode sequence (repeats of A,T and G), MspI site, primer priming site and homology arms, were synthesized by Integrated DNA Technology. The assembly reaction of 1:5 vector:insert ratio was carried out for 1 hour at 50C using NEBuilder HIFI DNA assembly kit (New England Biolabs, NEB). Assembly products were electroporated into NEB Turbo Competent E.Coli (NEB) and then plated on ampicillin-containing medium. Ampicillin resistant colonies were collected and extracted for plasmids using Maxiprep kit (Invitrogen). Ten sampling clones from the agar plate were analyzed by PCR and Sanger sequencing to verify successful cassette insertion and barcode diversity (Primer details in Table S1).

### Generation of founder cell library and cell lines

Barcoded reporter and third generation lentiviral packaging plasmids were transfected into HEK 293T cells to generate a library of barcoded lentivirus. Viral supernatant was collected and concentrated by Lenti-X-concentrator (Takara) at 48 hour post transfection. K562 cells were transduced with barcoded virus in cultured media supplement with 5 μg/ml polybrene and 20mM HEPES for 2 hours of spinoculation and 24 hours of incubation. An m.o.i of approximately 0.01, corresponding to 1% infectivity estimated by flow cytometer, was used to ensure that the majority of cells were labeled with single barcode per cell. Founder cells were selected by fluorescence-activated cell sorting (FACS) at 72 hours post transduction. Founder cells were expanded for two weeks and split into two pools. In the first pool, cells were subject to mapping the genomic location of barcoded reporter. In the second pool, cells were single-cell sort to establish cell lines of unique barcode.

### Identification of genuine barcode list

A library of genuine barcodes in founder cells was first listed. Briefly, barcode region was amplified in first nested PCR from 5 µg of genomic DNA in 50 ul of 20 cycle PCR reaction using Titanium Taq. Barcode amplicons were enriched from genomic DNA using SPRI beads(Beckman Coulter) and further amplified in the second nested PCR for 20 cycles. Illumina adapter was attached to final amplicon, amplified and sequenced on Illumina HiSeq 3000 platform (1×50bp). Sequencing reads were filtered and analyzed using Matlab Bioinformatics Toolbox. To identify genuine barcode, we used the following algorithm. First, we sorted barcodes according to their counts from most frequent to least frequent. Then, mutant versions of each barcode, defined as barcodes within a Hamming distance of 2, were sequentially removed. We consider remaining sequences as ‘‘genuine’’ barcodes. We recovered 756 genuine barcodes from 3,000 sorted founder cells.

### Mapping of reporter integration sites

Mapping of reporter integration sites was done by inverse PCR coupled with high-throughput sequencing. Briefly, founder cells were collected and splitted into two replicates. For each replica, 2 µg of genomic DNA was digested with 20 units of MspI (NEB) overnight at 37C in a volume of 100 µl. Subsequently, three sets of ligation reactions were set up by incubating 600 ng of purified digested DNA with 2 µl of high-concentration T4 DNA ligase (NEB, M0202T) overnight at 4C in a volume of 400 µl. The ligation reactions were purified by phenol-chloroform isoamyl alcohol extraction and ethanol precipitation. DNA pellets were dissolved in 30 µl of water. Two rounds of PCR were performed to amplify and enrich fragments containing both the barcodes and flanking genomic DNA regions (Primer details in Table S1). For the first round of nested PCR, five sets of 25-cycle reaction in a volume of 50 µl were performed using Phusion Hot Start Flex 2X Master Mix (NEB) and 5 µl of ligated products as templates. Amplicon was pooled together, cleaned by DNA Clean & Concentrator kit (Zymo), and diluted in 50 ul of water. For the second round of nested PCR, four sets of 15-cycle reaction in a volume of 50 ul were done with 5 ul of cleaned amplicon from first PCR. Purified sample was further ligated with Illumina adapter, amplified and sequenced on Illumina HiSeq 3000 platform (2×150bp). Sequencing reads were filtered and analyzed using Matlab Bioinformatics Toolbox. The genomic regions associated with genuine barcodes were extracted from mapping reads and aligned against the human genome (hg38) using STAR [85]. Detected integration sites from each replicate were compared and assigned to each genuine barcode only if top candidate site from both replicates are identical. Mapping of reporter integration sites were plotted on the ideogram (Fig.2b) using R and karyoploteR package [86]. Genome coordinate of reporter integration site was converted to human reference genome(hg19) using UCSC liftOver tool (Kuhn et al., 2013) for comparison to ChIP-Seq data.

### Combinatorial pool sequencing

Identity of reporter cell lines, linked by DNA barcodes, were simultaneously revealed in a single run using combinatorial pooled sequencing. Clonal numbers were encoded in a form of pooling pattern. To increase decoding accuracy, we designed pooling signature to be unique four selected pools out of total eighteen pools. Cells from each clone were splitted into four pools according to the design. Sequentially, genomic DNA from individual pool of mixed clones was extracted and used as templates for PCR to amplify barcode using same procedure described in the method of identification of genuine barcode list. Forward primers of second nested PCR contain 6-bp index DNA to label PCR products from each pool, which allow high-throughput multiplex sequencing (Primer details in Table S1). Sequences were filtered and demultiplexed using Matlab Bioinformatics Toolbox. Genuine barcodes from all pools were first listed. For each detected barcode, normalized counts per pools were calculated and pools showing high reads above the threshold were identified. Barcodes with four detected pools were first assigned to the clone showing matched pooling design. Some barcodes were found in more than four pools when sister cells, expanded from one founder cell, were sorted into multiple wells during single-cell sort. A list of merged pooling signature of two unassigned clones was matched with barcodes showing complexed readout. Clones with two inserted barcodes(∼2% of the population) were excluded from the library of reporter cell lines.

### Epigenetic drug treatment

We first created control cells expressing both mClover and IRFP670 from multiple integration sites. Briefly, K562 cells were transduced with CMV-IRFP670 lentivirus at high m.o.i. and sorted for IRFP positive cells. Lentiviral transduction of CMV-mClover was followed and dual reporter cells were selected by FACS. Control cells were co-cultured with individual reporter clones. 0.5 million cells of mixed samples were separately treated with DMSO, 400 nM of TSA, 5 µM of 5’ Azacytidine and 1 µM of JQ1 for 24 hours. Afterward, cells were collected and measured for expression distribution of mClover and IRFP using BD FACSCelesta flow cytometer. Experiments were done in three replicates per drug treatment per clone. IRFP expression was used to separate control cells from sample cells. To eliminate non loci-specific effects, log10-transformed mclover expression of control cells with epigenetic drug treatment was calibrated to match corresponding expression in DMSO condition. Same adjustment was applied to reporter cells in the same well. the Kolmogorov–Smirnov test was performed by Matlab software to compare histogram similarity of mClover distribution under different conditions.

### Validation of Combinatorial pool sequencing

For the validation of combinatorial pool sequencing, 5 clones were randomly chosen and two of them are sister clones, sharing same barcode. Genomic DNA (200ng) was used as a template for amplification with a set of validation primers (Primer details in Table S1). PCR products were cleaned by DNA Clean & Concentrator kit (Zymo) and Sanger sequenced to verify the barcode sequences.

### Analysis of ChIP-Seq data

Following ChIP-seq data sets were downloaded from ENCODE: H2A.Z (ENCFF191EXE), H3K4me1 (ENCFF526QTS), H3K4me2 (ENCFF118MMT), H3K4me3 (ENCFF715DGL), H3K9ac (ENCFF602QRW), H3K9me1 (ENCFF526UWC), H3K9me3 (ENCFF834YLI), H3K27ac (ENCFF010PHG), H3K27me3 (ENCFF445UCR), H3K36me3 (ENCFF678IWR), H3K79me2 (ENCFF003CLZ) and H4K20me1 (ENCFF143CUR). Fold change over control signals were averaged within a window of 10 kb centered around integration sites using R and Bioconductor packages.

### Methylation analysis of CMV promoter

To assess the levels of DNA methylation of CMV promoter, targeted bisulfite conversion was performed. Genomic DNA from clones of interest was extracted and bisulfite converted with Qiagen’s EpiTect bisulfite kit according to manufacturer’s instructions. Bisulfite converted genomic DNA was used as a template for two rounds of nested PCR using Invitrogen’s Phusion U polymerase (Primer details in Table S1). Amplicons from all samples were pooled together, purified by DNA Clean & Concentrator kit (Zymo) and deep sequenced. Sequencing reads were filtered and demultiplexed by barcode using Matlab Bioinformatics Toolbox. Changes in cytosine base at CpG sites were converted into a matrix of methylation status.

## Figure Legend

**Supplementary Figure 1:**
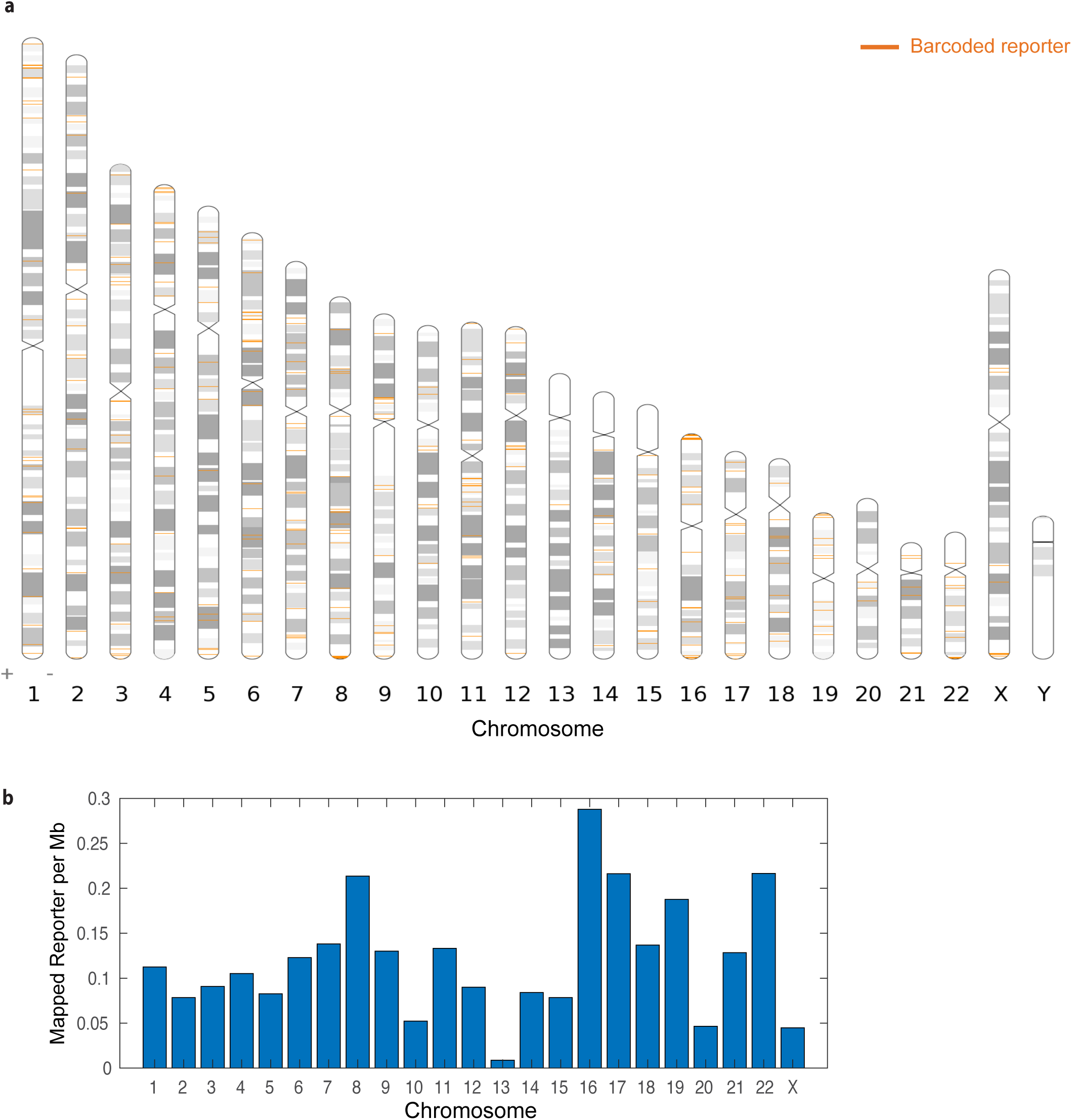
Genome-wide map of barcoded reporter insertion. **(a)** Positions of mapped barcoded-CMV-mClover reporters along all chromosome. Each synthetic reporter is represented as an orange tick on the ideogram. We detected 756 candidate genuine barcodes from three thousand original founder cells after two weeks of expansion. **(b)** A bar graph representing the frequency of mapped reporter per megabase.

**Supplementary Figure 2:**
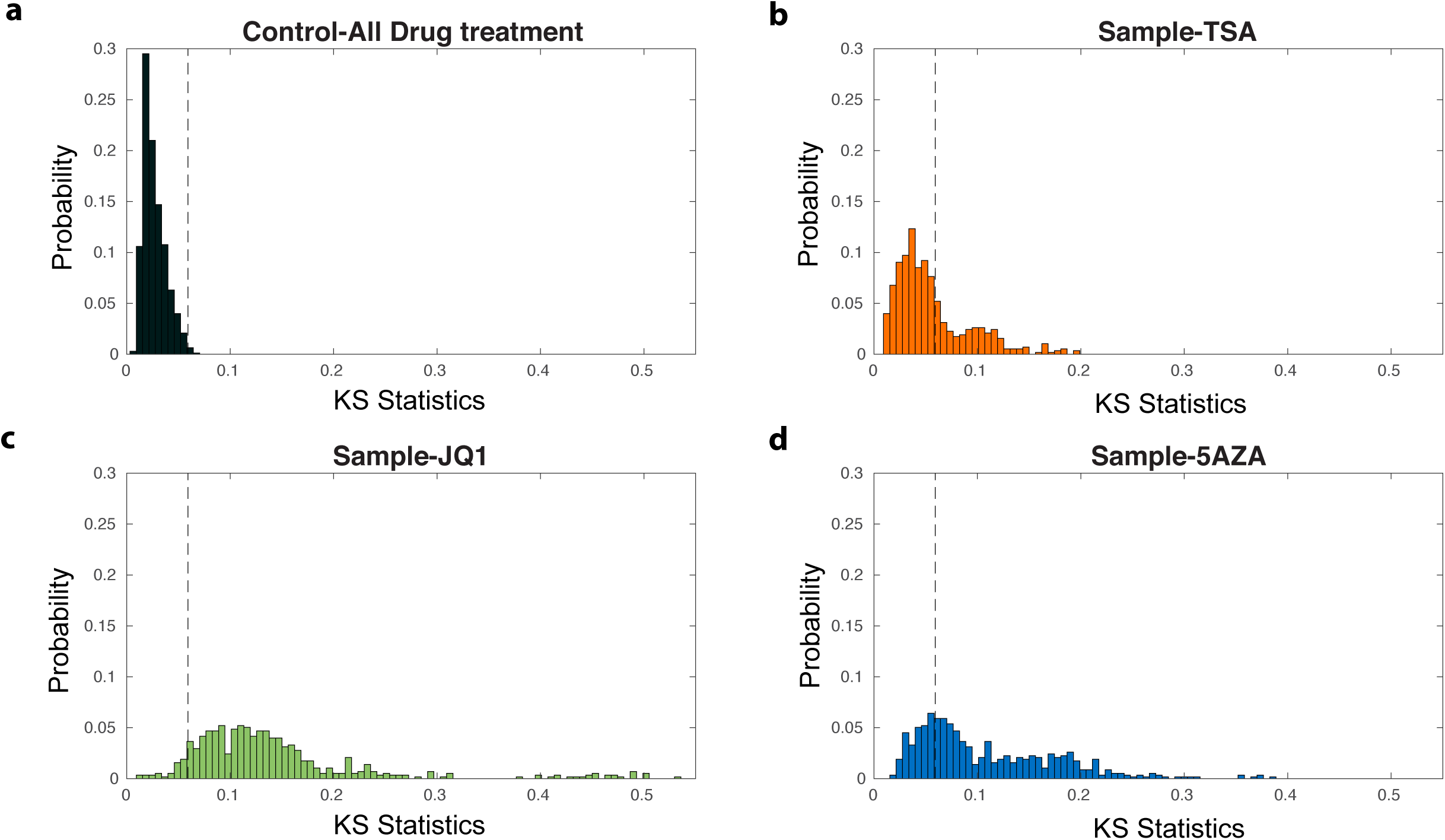
Drug screening shows different number of hits for different epi-drug treatment. **(a)** The distributions of KS statistics in control cells fall within the three standard deviation limits. **(b-d)** The number of hits in the drug screen for TSA(b), JQ1(c) and 5-AZA(d) is determined by Kolmogorov-Smirnov statistics passing three standard deviation limits of control groups.

